# Altering Polycomb Repressive Complex 2 activity partially suppresses *ddm1* mutant phenotypes in Arabidopsis

**DOI:** 10.1101/782219

**Authors:** Martin Rougée, Leandro Quadrana, Jérôme Zervudacki, Vincent Colot, Lionel Navarro, Angélique Deleris

## Abstract

In plants and mammals, DNA methylation is a hallmark of transposable element (TE) sequences that contributes to their epigenetic silencing. In contrast, histone H3 lysine 27 trimethylation (H3K27me3), which is deposited by the Polycomb Repressive Complex 2 (PRC2), is a hallmark of repressed genes. Nevertheless, there is a growing body of evidence for a functional interplay between these pathways. In particular, many TE sequences acquire H3K27me3 when they lose DNA methylation and it has been proposed that PRC2 can serve as a back-up silencing system for hypomethylated TEs. Here, we describe in the flowering plant *Arabidopsis thaliana* the gain of H3K27m3 at hundreds of TEs in the mutant *ddm1*, which is defective in the maintenance of DNA methylation specifically over TE and other repeat sequences. Importantly, we show that this gain essentially depends on CURLY LEAF (CLF), which is one of two otherwise partially redundant H3K27 methyltransferases active in vegetative tissues. Finally, our results challenge the notion that PRC2 can be a compensatory silencing system for hypomethylated TEs, as the complete loss of H3K27me3 in *ddm1 clf* double mutant plants was not associated with further reactivation of TE expression nor with a burst of transposition. Instead, and surprisingly, *ddm1 clf* plants exhibited less activated TEs, and a chromatin recompaction as well as hypermethylation of linker DNA compared to *ddm1*. Thus, we have described an unexpected genetic interaction between DNA methylation and Polycomb silencing pathways, where a mutation in PRC2 does not aggravate the molecular phenotypes linked to TE hypomethylation in *ddm1* but instead partially suppresses them.

**Author summary:** Epigenetic marks are covalent modifications of the DNA or its associated proteins (Histones) that impact gene expression in a heritable manner without changing DNA sequence. In plants and mammals, DNA methylation and trimethylation of Lysine 27 of Histone 3 (H3K27me3) are two conserved, major epigenetic systems that mediate the transcriptional silencing of transposons (invasive mobile genetic elements) and of developmental genes respectively. However, in the absence of DNA methylation, H3K27me3 marks can be recruited to transposons, suggesting that the two silencing systems can be compensatory. To test this hypothesis, we analyzed a compound DNA methylation and H3K27me3 mutant of the plant model *Arabidopsis thaliana* (importantly, mammals harboring equivalent mutations would not be viable). First, this approach allowed us to gain mechanistic insights into the recruitment of H3K27me3 at transposons. Furthermore, we also showed that transposon silencing release in the DNA methylation mutant was not enhanced, contrary to our initial hypothesis, but, surprisingly, partially suppressed by a mutation in H3K27me3 deposition. Thus, our genomic analysis revealed an unexpected and antagonistic genetic interaction between two major silencing pathways whose interplay is at the heart of many biological processes, including cancer.

## Introduction

DNA methylation is an epigenetic mark involved in the stable silencing of transposable elements (TEs) as well as the regulation of gene expression in plants and mammals. When present over TE sequences, it is usually associated with the di- or trimethylation of histone H3 lysine 9 (H3K9me2/H3K9me3) and, in plants, positive feedback loops between the two marks exist, which maintain TE sequences in the heterochromatic state (1,2). In *A. thaliana*, DOMAINS REARRANGED METHYLASE 2 (DRM2) establishes DNA methylation in all three sequences contexts (i.e. CG, CHG and CHH) in a pathway referred to as RNA-directed DNA methylation (RdDM) that involves small RNAs (3,4). Maintenance of DNA methylation over TEs is achieved by the combined and context-specific action of RdDM (CHH methylation), CHROMOMETHYLASES 2 and 3 (CMT2 and CMT3, for CHH and CHG methylation, respectively) (5,6) and METHYLTRANSFERASE1 (MET1) (CG methylation) (7). In addition, the SNF2 family chromatin remodeler DECREASE IN DNA METHYLATION 1 (DDM1) is necessary for DNA methylation of most TE sequences in all cytosine sequence contexts.

In contrast to DNA methylation, histone H3 lysine 27 trimethylation (H3K27me3), which is targeted by the highly conserved Polycomb Group (PcG) proteins, in particular Polycomb Repressive Complex 2 (PRC2), is a hallmark of transcriptional repression of protein-coding and microRNA genes in plants as well as in animals (8–11); it is thought to act by promoting a local compaction of the chromatin that antagonizes the transcription machinery (11) in order to maintain transcriptional silencing (12). Thus, H3K27me3 and DNA methylation are generally considered as mutually exclusive chromatin marks.

Nonetheless, there is a growing body of evidence of an interplay between the two silencing pathways. In particular, in both plants and mammals, many TE sequences gain H3K27me3 upon their loss of DNA methylation (13–16). Moreover, in the filamentous Neurospora, H3K27me3 is redistributed from gene to TE-rich constitutive heterochromatin when the heterochromatic mark H3K9me3 or the protein complexes that bind to it are lost (17).

The fact that H3K27me3 can mark TE sequences upon their demethylation led to the idea that PcG could serve as a back-up silencing system for hypomethylated TEs (14). Consistent with this notion, subsequent work in mammals showed that H3K27me3 re-established the repression of thousands of hypomethylated TEs in embryonic stem cells subjected to rapid and extensive DNA demethylation (18).

In a previous study, we provided evidence supporting a role of PcG in the transcriptional silencing of *EVD* (19), an *A. thaliana* retroelement of the *ATCOPIA93* family that is tightly controlled by DNA methylation and which transposes in plants mutated for the chromatin remodeler *DDM1* (20). We observed that silencing of *EVD* is dependent on both DNA methylation and H3K27me3, which, at this locus, depends on the SET-domain protein CURLY LEAF (CLF) (19). Whether the dual control observed at *EVD* is also present at other plant TEs is unknown.

In the present work, we integrated genetics, epigenomics and cell imaging to show that numerous TEs gain H3K27me3 in response to *ddm1*-induced loss of DNA methylation. We demonstrate also that this gain is mediated by CLF, with no apparent role for the SET-domain H3K27 methyltransferase SWINGER (SWN), which is also active in vegetative tissues and otherwise partially redundant with CLF (21–23). Unexpectedly, the combination of *ddm1* and *clf* mutations was not associated with further reactivation of TE expression or transposition as compared to *ddm1*, except for *EVD*. Instead less TEs were expressed and we observed a partial recompaction of heterochromatin coupled with DNA hypermethylation, prominently in linker DNA, in *ddm1 clf* versus *ddm1*. Together, these results show that altering PRC2 activity partially suppresses *ddm1* phenotypes.

## Results

### Hundreds of hypomethylated TEs gain H3K27me3 in *ddm1*

We previously showed that in *met1* mutants impaired for CG methylation, hundreds of TEs gain H3K27me3 methylation in Arabidopsis (14). While *met1* and *ddm1* mutants are both globally hypomethylated, the later, in contrast to the former, affects almost exclusively TE and other repeat sequences, and in the three sequence contexts. Thus, the *ddm1* mutant appeared to be a more relevant background to directly explore the interplay between DNA methylation and Polycomb at TE sequences and we conducted a H3K27me3 ChIP-seq experiment in this mutant. Like in *met1*, hundreds of TEs showed increased accumulation of H3K27me3 in *ddm1* (**Fig 1A-C**). A subset of 672 TEs showed no H3K27me3 ChIP-seq signal in wild-type plants and significantly gained H3K27me3 over their full length in *ddm1* (**Fig 1D**). Moreover, the vast majority of those 672 TEs are located in pericentromeric regions (**Fig 1E**) and were included in the subset of TEs that gain H3K27me3 in *met1* (14) (**S1A Fig**). Two major TE super families were overrepresented among the 672 TEs (LTR/Gypsy, DNA/others) as compared to the distribution of the heterochromatic, pericentromeric TEs (targets of DDM1) families (**S1B Fig**), suggesting the existence of sequence-specific targeting. Indeed, PRC2 can be targeted to specific genes by the recognition of short sequence motifs (24). Consistent with this motif-based targeting, we detected a significant enrichment of the Telobox, CTCC, GA-repeat and AC-rich motifs in the 672 TEs compared to the rest of the heterochromatic TEs (**Fig 1F-G**). These results suggest the presence of an instructive mechanism of PRC2 recruitment at TEs with particular motifs used as nucleation sites. In addition, the detection of continuous blocks of H3K27me3, in particular on chromosomes 1 and 4 (**S1C Fig**), may suggest that, once nucleated, H3K27me3 domains spread over entire TE sequences and even beyond, into nearby TEs, in a fashion similar to what was described at genic sequences (22,23). Of note, redistribution of H3K27me3 to TEs in *ddm1* did not seem associated with a loss of H3K27me3 at genes (0 gene lost H3K27me3 significantly in *ddm1*) contrary to what was observed in *met1* (14), presumably because less TEs become targets of PRC2 in *ddm1* versus *met1*.

**Figure 1.**
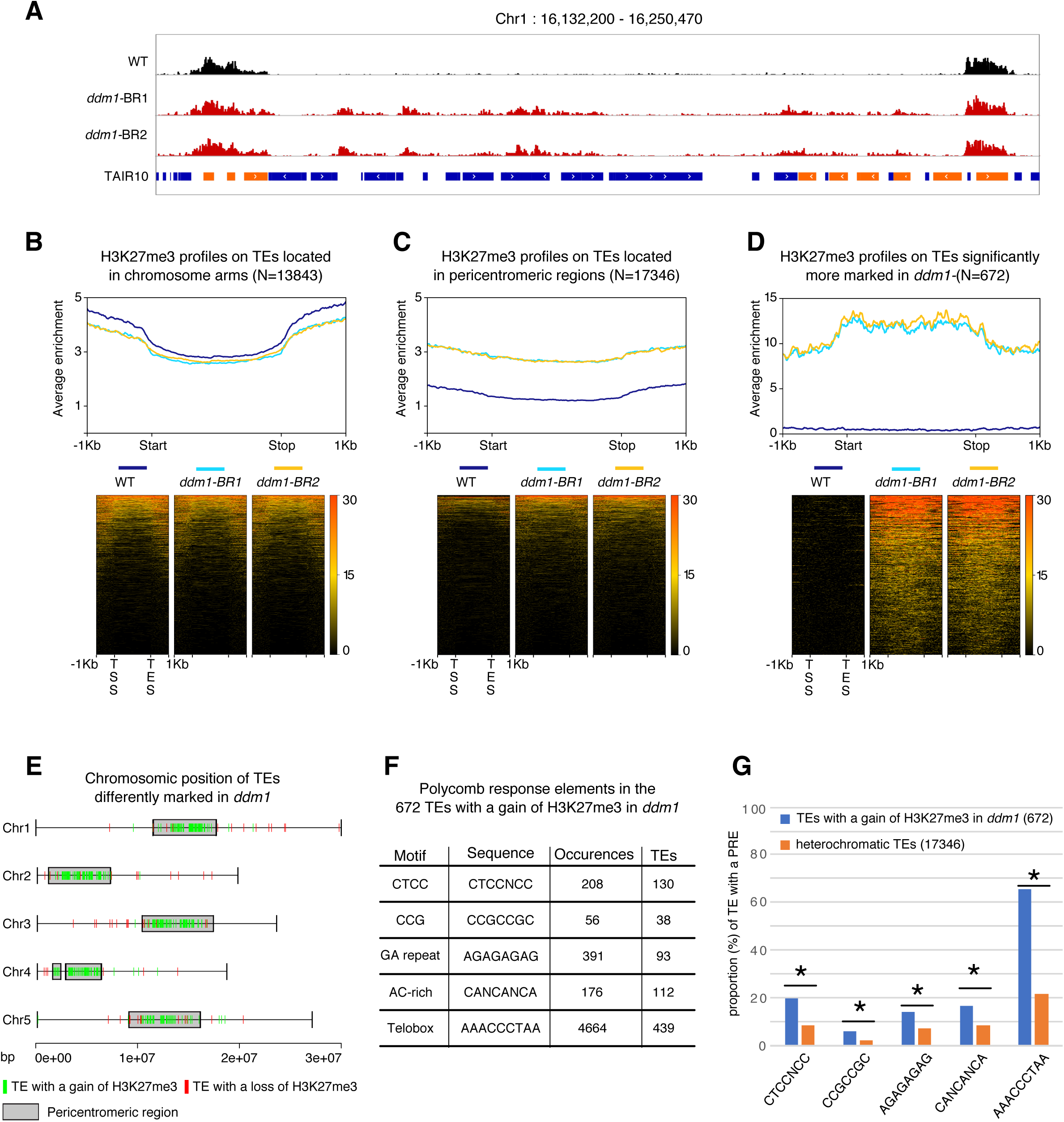
A mutation in *DDM1* leads to a gain of H3K27me3 on some heterochromatic TEs. **(A)** Representative genome browser view of H3K27me3 levels in a 100 kb heterochromatic region of chromosome 1 in wildtype (WT) and two biological replicates (BR) of *ddm1* (blue bar: TE; orange bar: gene). **(B**,**C**,**D)** H3K27me3 enrichment in WT, *ddm1*-BR1 and *ddm1*-BR2 over TEs located in euchromatin (B), heterochromatin **(C)** and TEs with a significant positive fold change (FC) of H3K27me3 in *ddm1* (Pval<0,1; Log 2FC>2) **(D)**. The upper graph shows the mean enrichment of H3K27me3 over a given TE subset; the corresponding heatmap below ranks them from top to bottom according to the average enrichment in all genotypes. **(E)** Chromosomal distribution of the 672 TEs that significantly gain H3K27me3 (Pval<0,1; Log2FC>2) and the 61 TEs that significantly lose H3K27me3 in *ddm1* (Pval<0,1; Log2FC<-2). **(F)** Table showing the occurences of Polycomb Response Elements (PRE) (Xiao et al., 2017) among the 672 TEs with a gain of H3K27me3 in *ddm1*. **(G)** Graph showing the proportion of TEs (TEs with a gain of H3K27me3 in *ddm1* compared to heterochromatic TEs) with a given PRE (* indicates a significant difference Pval<0.5, t-test).

### The gain of H3K7me3 over hypomethylated TEs depends on CLF

To test whether CLF is required for the gain of H3K7me3 over TEs we performed a second set of ChIP-seq experiments in two different F3 progenies of *ddm1 clf* double mutant (and that we refer to as Biological Replicates, here BR1 and 2). Because total levels of H3K27me3 are strongly reduced in both *clf* and *ddm1 clf* mutants (**S2A Fig**) compared to wild-type, we spiked-in exogenous Drosophila chromatin in the chromatin extracts for normalization (25). There was no consistent difference in the levels of H3K27me3 at genes between *ddm1 clf* and *clf* mutant (**S2B Fig**), in accordance with DDM1 affecting TE and other repeat sequences specifically (26). Conversely, the gain of H3K27me3 observed over TEs in *ddm1* was almost completely abolished in *ddm1 clf* (**Fig 2A-C and S2C Fig**). Together, these results show that deposition of H3K27me3 at most TEs in *ddm1* is fully dependent on *CLF* with no apparent role of the paralogous histone methyltransferase *SWN*. This is in contrast with the well-established, partial dependency of H3K27me3 deposition at genes on CLF (22,23), which has been proposed to be due to a specialization of this factor in mediating amplification and spreading of H3K27me3 marks after their establishment by SWN (23). Thus, at least over *ddm1*-dependent hypomethylated TEs, CLF is required for the nucleation, and perhaps, additionally, spreading of H3K27me3.

**Figure 2.**
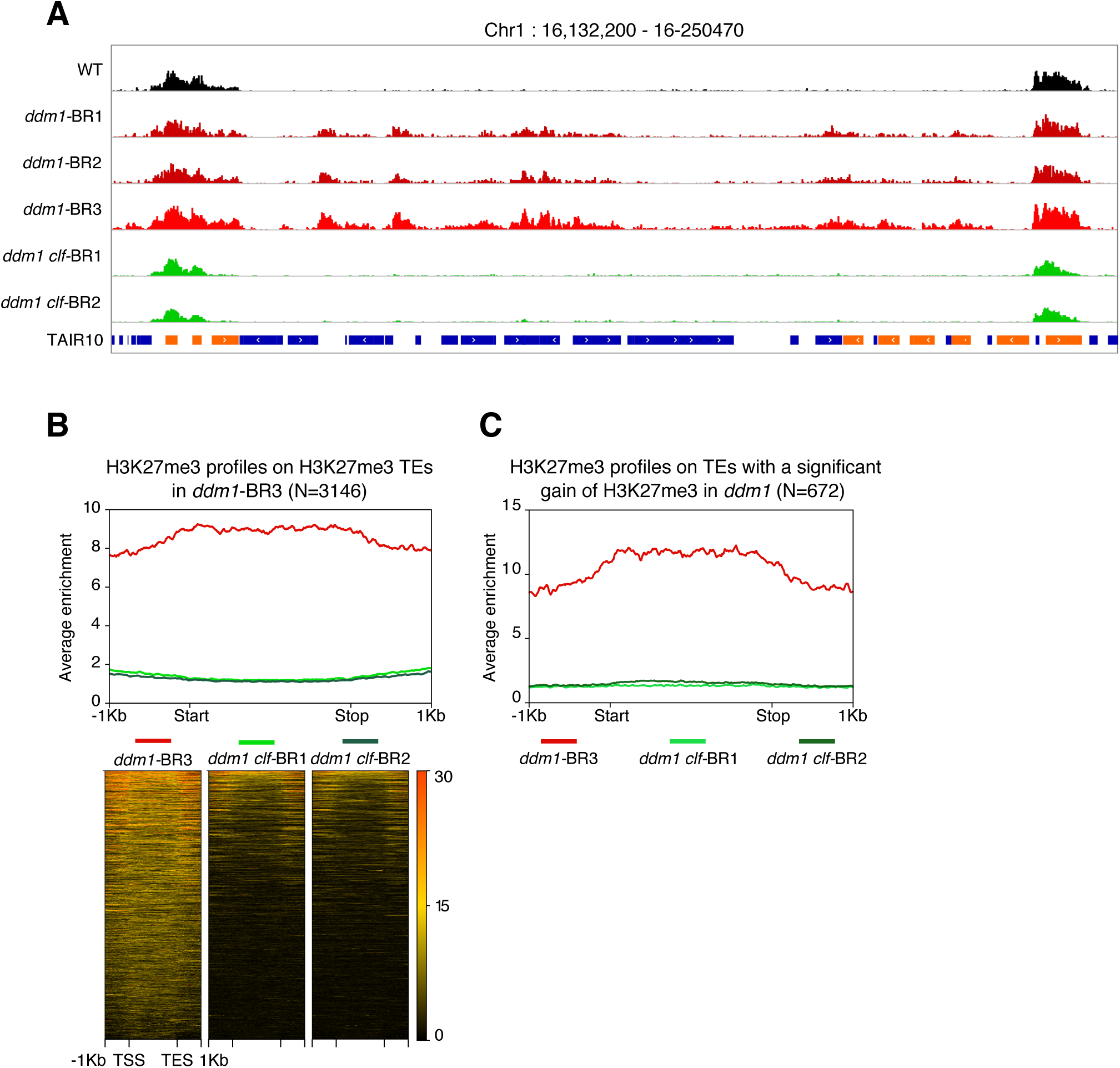
The gain of H3K27me3 in *ddm1* is lost in *ddm1 clf*. **(A)** Genome browser view of H3K27me3 levels on the same region as Figure 1 **(B)** H3K27me3 enrichment in *ddm1* and two *ddm1 clf* replicates over TEs with a H3K27me3 domain in *ddm1* with corresponding heatmap ranking TEs according to the H3K27me3 mean value. **(C)** Metagene representing H3K27me3 enrichment in *ddm1*-BR3, *ddm1 clf*-BR1 and *ddm1 clf*-BR2 of the 672 TEs previously shown to gain H3K27me3 in *ddm1*.

### TE activation in *ddm1* is not enhanced in *ddm1 clf*

We previously described the phenotype of *ddm1 clf* mutants at the rosette stage (4.5 weeks-old) in F2 plants (19). In all mutant lines, we further observed floral defects (**S3A Fig**) which severity seems to increase with generation time, resulting in almost complete sterility of F4 generation mutants. In addition, by examining the plants at the seedling stage (15 days-old), we could notice additional phenotypes in two out of three independent *ddm1 clf* lines (chlorosis in one line, growth arrest in another) that segregated 1:3 and thus evoked the segregation of recessive mutations (**S3B Fig**). We previously showed that there is an enhanced accumulation of *ATCOPIA93* mRNAs in *ddm1 clf* double mutant as compared to *ddm1* (19), thus we hypothesized that these segregating phenotypes were caused by the transposition of TEs activated in *ddm1 clf*.

To test whether *ATCOPIA93* activity is increased and whether additional copies of this TE family accumulate in *ddm1 clf*, we performed Southern blot analysis (**Fig 3A**) using a probe specific of *ATCOPIA93 EVD*/*ATR* (*ATR* is an almost identical copy of *EVD*). We could only observe one copy of *EVD* and *ATR* in the wild-type, *clf* single mutants as well as in the *ddm1* 2nd generation inbred mutants. This result was not surprising since *EVD* was previously found to be active in *ddm1* or *ddm1*-derived epigenetically recombinant inbred lines but this was observed in late generations (8_th_ and beyond) (20,27). By contrast, we observed linear extrachromosomal DNA in all the *ddm1 clf* double mutant lines tested (progenies of individual F2), consistent with previous results (19); in addition, in two out of three double mutant lines, we detected numerous additional bands corresponding to *EVD*/*ATR* new insertions (**Fig 3A**). This indicates that transposition occurs in the double mutant, although to various extents which may reflect different dynamics of mobilization following the initial event (27). Pyrosequencing performed on the mutants showed that *ATCOPIA93* DNA (extrachromosomal and integrated) was mostly contributed by *EVD* (**Fig 3B**). Thus, DNA methylation and H3K27me3 act in synergy to negatively control *EVD* activity and prevent its transposition, in accordance with the dual control they exert on its transcription (19). To test whether more TEs are activated in *ddm1 clf*, we performed TE-sequence capture (28,29), which enables sensitive detection of new TE insertions at the genome wide-level. Essentially, the capture probes cover 200bp at each end of 310 potentially mobile TEs, that belong to 181 TE families either identified as mobile in various Arabidopsis ecotypes using the split-read approach, or for which non-degenerate and thus potentially mobile copies are present in the Col-0 genome (29). Our rationale was that the activity of some of these TEs is weak or absent in the Col-0 *ddm1* mutants or their derived epigenetic recombinant inbred lines because there is a second layer of silencing mediated by H3K27me3 in these epigenetic mutant backgrounds. Using about 100 seedlings of wild-type, *ddm1* (2nd generation inbred mutants) and *ddm1 clf* (F3 progenies), no new insertions were detected for any of the 250 TEs tested in this manner, except for *EVD*, which had accumulated more copies in *ddm1 clf* than in *ddm1* (**Fig 3C)**, although this difference was not statistically significant in average — this was presumably because the *ddm1 clf* line (#1), where only ecDNA accumulates, was included in the experiment along lines #2 and #3 **(S3C Fig**).

**Figure 3.**
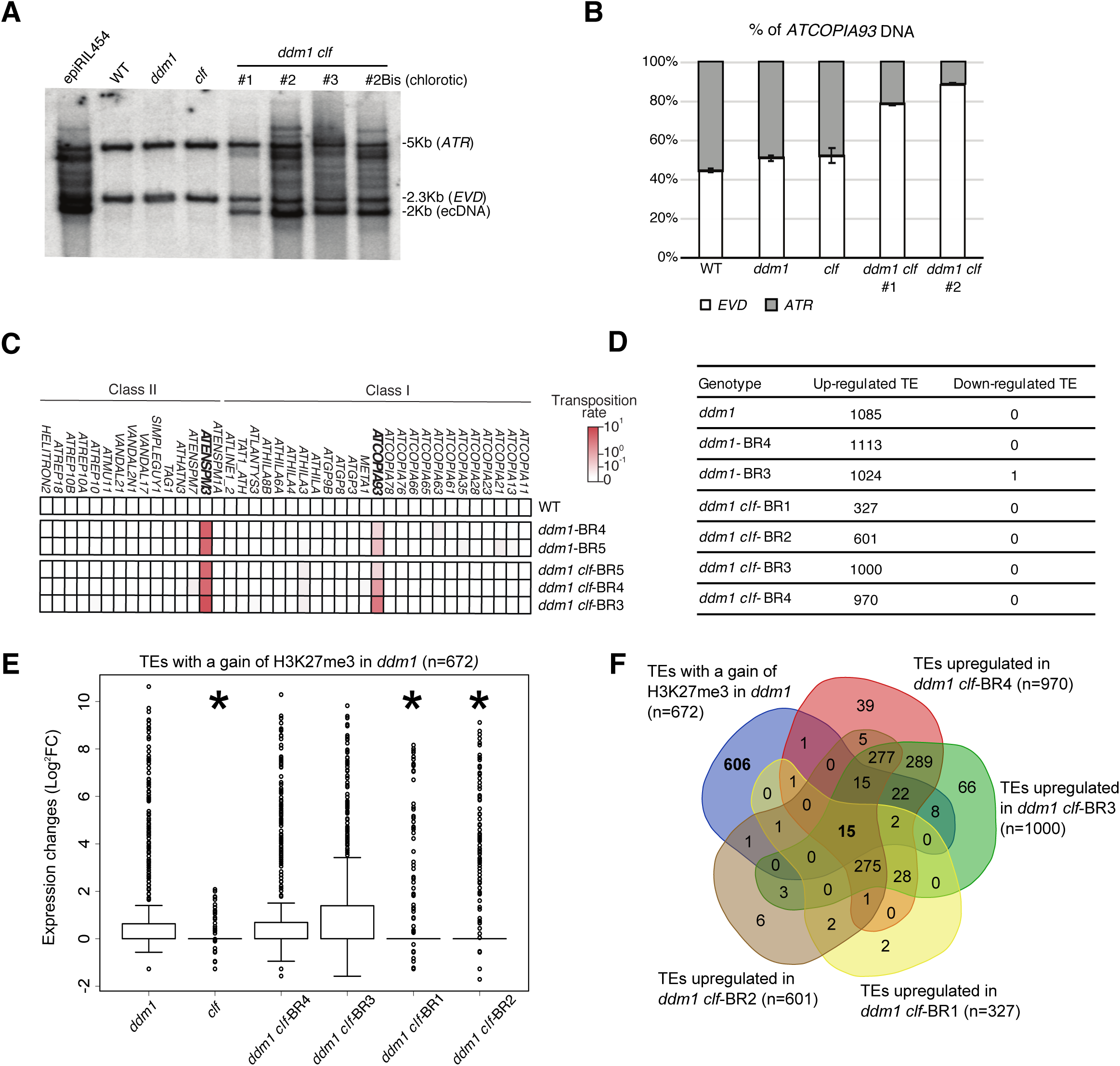
Transposons activation in *ddm1* and *ddm1 clf*. **(A)** Southern blot analysis on a pool of leaves (8 rosettes for each genotype), using an *ATCOPIA93* probe that recognizes both *EVD* and *ATR* copies; epiRIL454 (Mari-Ordonez et al., 2013) was used as a positive control for activation of *ATCOPIA93* and three independent *ddm1 clf* lines were tested. **(B)** Qualitative analysis of genomic DNA by pyrosequencing. The position interrogated corresponds to the discriminating SNP between *EVD* (C/G) and *ATR* (A/T) and the % indicated represent the % of G (*EVD*, white bar) or T (*ATR*, grey bar) **(C)** Heatmap showing the transposition rates of 40 potentially mobile TE families as determined by TE-capture (Quadrana et al., 2016) using 100 seedlings for each genotype. **(D)** Table showing up-regulated and down-regulated TEs compared to WT as identified by RNA-seq analyses on seedlings. **(E)** Box plot of log2 RNA fold changes (mutant/WT) over the TEs that significantly gain H3K27me3 in *ddm1;* * indicates a significant decrease in the fold change for a given *ddm1 clf* line relative to *ddm1* (Pval<0,05, t-test). **(F)** Venn diagram showing the overlap between TEs with a significant gain of H3K27me3 in *ddm1* and TEs upregulated in the 4 different *ddm1 clf* lines.

To investigate further the relationship between H3K27me3 and TE silencing, we performed RNA-seq experiments in three and four biological replicates (BR) of *ddm1* and *ddm1 clf* (four different F3 progenies), respectively. In all *ddm1* populations, about a thousand TEs (929) were upregulated, the large majority of which (865) did not show a significant gain of H3K27me3 marks in this background. We found that a similar (*ddm1 clf*-BR3/4) or even lower (*ddm1 clf*-BR1/2) number of TEs were transcriptionally active in *ddm1 clf* compared to WT (**Fig 3D**) in keeping with the lack of enhanced transposition in *ddm1 clf*. In addition, these transcriptionally active TEs were expressed at similar levels (*ddm1 clf*-BR3/4) or less expressed (*ddm1 clf*-BR1/2) in *ddm1 clf* than in *ddm1* (**S3D Fig**). As for the TEs with a significant gain of H3K27me3 in *ddm1* (N= 672), we did not observe that they were more expressed upon complete loss of H3K27me3 in *ddm1 clf* than in *ddm1* : rather, they tended to be even less expressed than in *ddm1* in two *ddm1 clf* progenies (BR1/2) out of four (**Fig 3E**). Accordingly, among these TEs with a significant gain of H3K27me3 in *ddm1*, we found only 15 TEs that were transcriptionally active in all *ddm1 clf* F3 progenies (**Fig 3F)**. Thus, with the exception of *EVD* case study, which supports our initial hypothesis of a dual epigenetic control by DNA methylation and H3K27me3, it appears that *ddm1*-induced activation of TEs is not enhanced in *ddm1 clf* and, rather, tends to be partially suppressed in this double mutant background, with some heterogeneity between the lines used.

### *ddm1 clf* displays chromatin recompaction and increased DNA methylation compared to *ddm1*

To understand the mechanisms underlying the antagonism between DNA methylation and *CLF*-dependent H3K27me3 deposition, we performed cytogenetic analyses on nuclei from single and double mutants for *DDM1* and *CLF*. Chromocenters formation and DNA compaction were unchanged in *clf* but strongly impacted in *ddm1* as previously published (30). By contrast, they were only partially affected in the double mutant, indicating that H3K27me3 loss induces chromatin recompaction specifically in the *ddm1* background (**Fig 4A-B**). In addition, immunofluorescence experiments showed that this chromatin recompaction was associated with the recovery of H3K9me2 distribution (**Fig 4C**).

**Figure 4.**
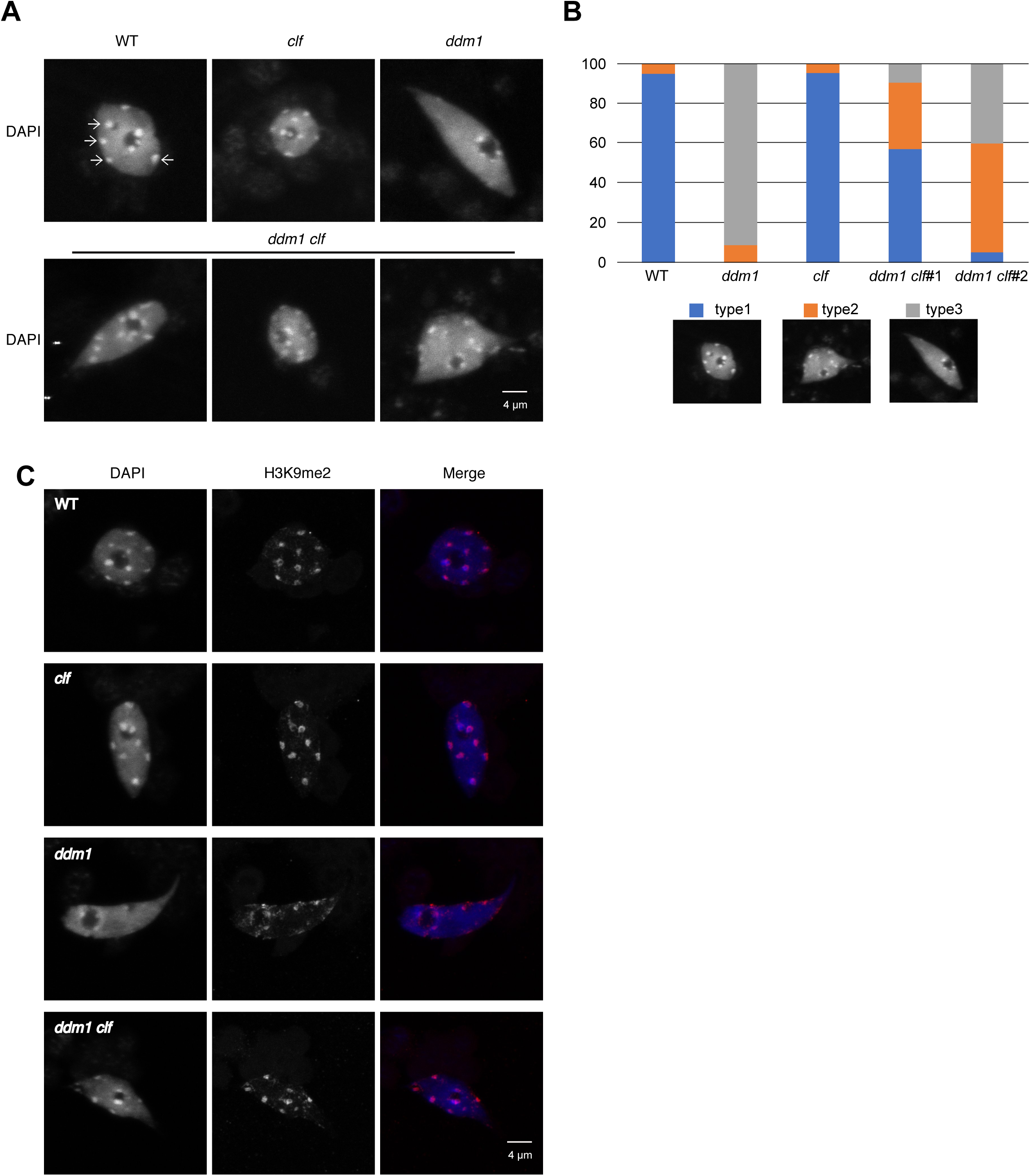
The double mutant *ddm1 clf* displays a partial recompaction of DNA and more visible H3K9me2 compared to *ddm1*. **(A)** Representative pictures of DAPI stained nuclei extracted from 50 fixed 10 days-old Arabidopsis seedlings for each genotype (three independent F3 lines from *ddm1 clf*). **(B)** Nucleus types were categorized according to the signal distribution relative to the heterochromatic chromocenters. Type1 compacted (like WT), type2; semi-compacted and type 3; decompacted (like *ddm1*). Data are presented as percentage and were derived from pictures of at least 20 nuclei from 3 independent experiments and two *ddm1 clf* lines **(C)** Representative pictures of DAPI staining and H3K9me2 immunodetection of nuclei extracted from 50 fixed 10 days-old Arabidopsis seedlings for each genotype.

In order to assess whether chromatin recompaction associates with DNA methylation gain, we compared the DNA methylomes of *ddm1* and *ddm1 clf* mutant plants. Consistent with the cytogenetic analyses, hundreds of loci exhibited higher DNA methylation in all three sequence contexts in *ddm1 clf* compared to *ddm1* (**Fig 5A**), with gains being generally most pronounced at CHH sites and variable from one line to the other (**Fig 5B**).

**Figure 5.**
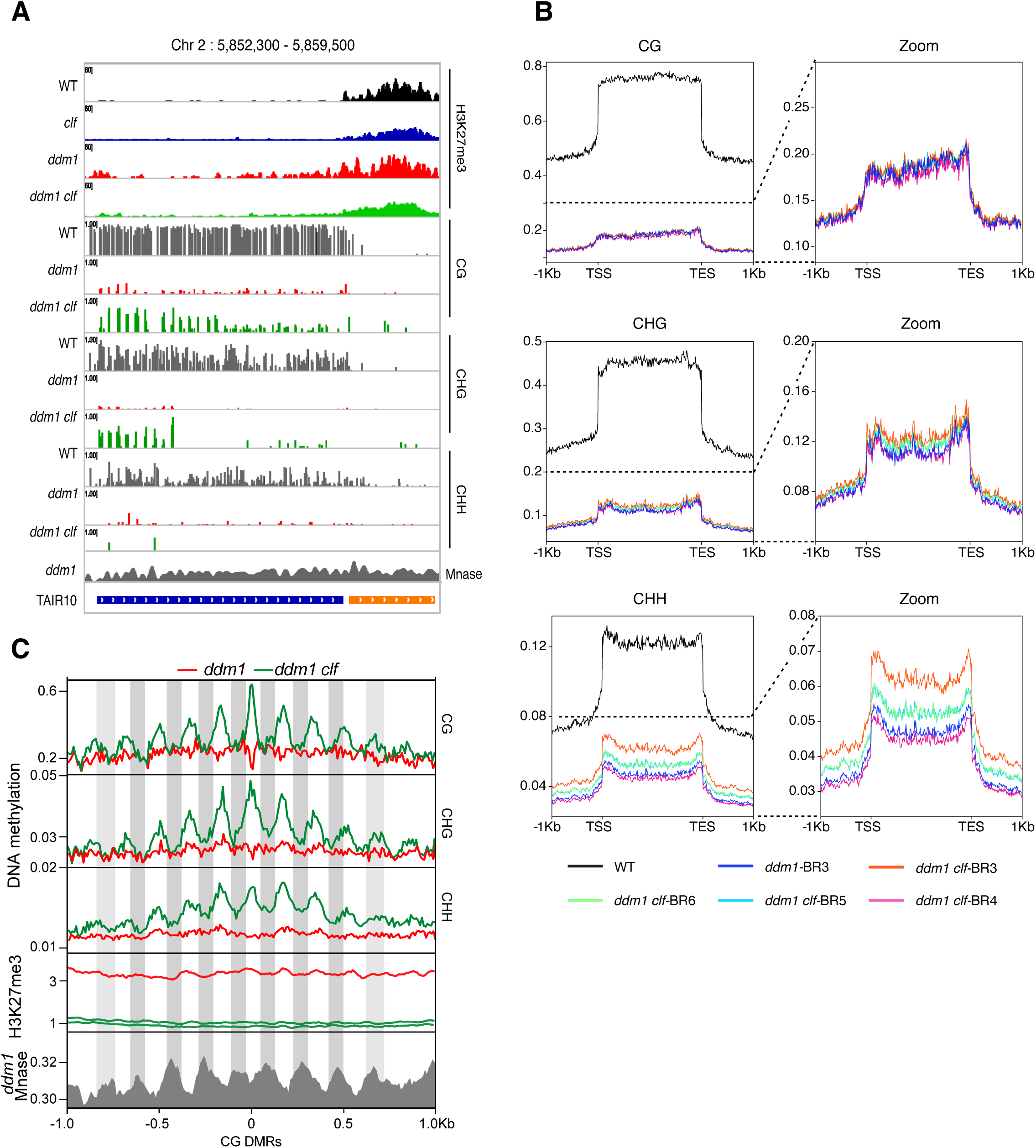
Periodic hypermethylation is observed in *ddm1 clf* compared to *ddm1*. **(A)** Representative genome browser view of H3K27me3 and methylation of a TE (blue bar) showing hypermethylation in *ddm1 clf* compared to *ddm1*. **(B)** Meta-TEs showing average DNA methylation levels on all TEs, in the three different contexts, in all the genotypes of interest, with a zoom on *ddm1* and *ddm1 clf* methylated TEs. **(C)** Plot showing average DNA methylation, H3K27me3 and nucleosome occupancy (every peak in *ddm1* Mnase data shows the position of a nucleosome) 1Kb upstream ans downstream of “small” CG DMRs (20bp) in the three different sequence contexts.

To test whether H3K27me3 directly antagonizes DNA remethylation in *ddm1* (or promotes *ddm1*-induced DNA methylation loss) in *cis*, we searched for differential methylated regions (DMRs) using non-overlapping 100bp windows. There were few hyper DMRs and these tended to be inconsistent across replicates, suggesting that they were the result of stochastic variation of DNA methylation (31). Moreover, TEs showing clear DNA methylation gain in *ddm1 clf* compared to *ddm1* (**Fig 5A**) were not identified by our DMR detection. Visual inspection of these DNA hypermethylated TEs revealed that increased DNA methylation was in fact confined to short sequences (∼20bp), typically separated by ∼140 bp (**Fig 5B, S5A Fig**). Indeed, we identified 1208 TE-specific short (20bp) sequences with higher CG methylation in *ddm1 clf* compared to *ddm1*. These small-size hyper-DMRs were distributed over 759 TEs, most of which (N= 571) are pericentromeric, and they were associated with a reduction of H3K27me3 levels (**Fig 5C**, H3K27me3 profiles centered on CG-DMRs). Among these 759 TEs hypermethylated in *ddm1 clf* compared to *ddm1*, 61 showed a significant gain of H3K27me3 in *ddm1:* this is a significant 3.8 representation factor enrichment (p<3.644e_-19_) compared to the number of TEs in the genome that would be both CG-hypermethylated (in *ddm1 clf*) and H3K27me3-marked (in *ddm1*) by chance. Although significant, this overlap is small, which suggests that globally the antagonism between H3K27me3 and DNA methylation is indirect. This could be explained by the impact of *ddm1 clf* double mutation on the transcriptome; nevertheless we did not find any consistent expression changes for the components involved in DNA methylation (**S5B Fig**).

Finally, the ∼140 bp distance between DMRs at discrete loci —which is roughly the size of nucleosomal DNA— prompted us to test whether DNA hypermethylation in *ddm1 clf* versus *ddm1* occurred preferentially between nucleosomes. Analysis of publicly available MNase data obtained for *ddm1* (32) indicated that the periodic DNA hypermethylation in *ddm1 clf* associate with linker DNA (**Fig 5C)**, consistent with *in vitro* and *in vivo* data showing that inter-nucleosomal DNA is more accessible to DNA methyltransferases than DNA bound to the nucleosome core particle (32). These findings suggest therefore that CLF-dependent H3K27me3 marks, although indirectly, limit the accessibility of DNA methyltransferases to linker DNA.

## Discussion

In this study, we have shown that hundreds of TEs gain H3K27me3 when hypomethylated by *ddm1*. This phenomenon has two important mechanistic implications, the first one being that DNA methylation can globally exclude H3K27me3. We and others previously showed that loss of DNA methylation induced by another hypomethylation mutant, *met1*, rather than loss of H3K9me2, allows for H3K27me3 deposition at TEs (13,14). However, another commonality between *met1* and *ddm1* mutants is the decompaction of their DNA, which could also contribute to favor the access of PRC2 to chromatin, a possibility that could be further investigated by identifying and using genetic backgrounds impaired for chromatin compaction but not DNA methylation. The second mechanistic implication of our observations is that PRC2 can be recruited to TEs. Here, we have characterized the transposons marked by H3K27me3 in *ddm1* and found some motifs recently described in the genic targets of PRC2 and functionally involved in the recruitment of this complex at genes (24). Thus, while some TEs could be marked by H3K27me3 due to spreading from adjacent PcG targets as it is the case for *EVD* (19), the presence of these motifs as well as the localization of many TEs far away from any H3K27me3 domain, suggests a sequence-based, instructive mode of *cis*-recruitment for PRC2. Alternatively, this recruitment could also be promoted by the enrichment of H2A.Z over demethylated TEs (33) since a mechanistic link was found between H2A.Z and H3K27me3 (34). These non-mutually exclusive scenarios will deserve future investigation.

Subsequently, the compound mutant of DNA methylation and PRC2, *ddm1 clf*, uncovered unexpected and interesting molecular phenotypes. First, PRC2 targeting of TEs in the *ddm1* background depends almost completely on CLF. This requirement of CLF contrasts with the partial dependency on CLF and SWN at H3K27me3 marked genes, where SWN plays a major role, presumably in nucleating H3K27me3 (23). A possible explanation for this result could be that the transcription factors implicated in the nucleation of H3K27me3 (24) at TEs in *ddm1* recruit a FIE-EMF2-PRC2 complex that does not contain SWN. Secondly, from a functional point of view and rather unexpectedly, our results did not show that PRC2 acts as a back-up silencing system in the absence of DNA methylation. Rather, they are reminiscent of the observations made in Neurospora where the gain of H3K27me3 in constitutive heterochromatin domains is not compensatory for gene silencing upon loss of H3K9 methylation (or its reader HP1) but rather leads to growth defects and altered sensitivity to genotoxic agents (17). In our study, in *ddm1 clf*, the loss of H3K27me3 did not enhance the number of *ddm1*-activated TEs. Instead, molecular phenotypes of partial *ddm1* suppression were observed: same or lesser number of activated TEs (similarly activated as in *ddm1* or even less) and partial compaction of DNA compared to *ddm1* associated with hypermethylation of the DNA in the three sequence contexts. Since the hypermethylation also occurred at TEs that are non-targets of PcG, we could not conclude on a direct antagonism between H3K27me3 and DNA methylation. Instead, we propose two possible mechanisms for the global hypermethylation of *ddm1 clf* compared to *ddm1*. Firstly, it is likely that in *ddm1* mutant, like in *met1* (33), H2AZ is incorporated at TEs. H2AZ is known to be antagonistic to DNA methylation and given that a mutation of *CLF* leads to loss of H2AZ (34), this may favor the reestablishment of DNA methylation. Another non-exclusive possibility could be that the chromatin decompaction displayed by *ddm1* in the presence of PRC2 globally restricts DNA remethylation which would occur upon recompaction in *ddm1 clf* through unknown mecanisms. In addition, small RNAs could also participate in either of these processes. What favors DNA compaction upon loss of H3K27me3, and, accordingly what antagonizes chromocenters formation in *ddm1* in the presence of H3K27me3 is also unclear. It could be linked to the replacement of H3.1, the substrate for ATRX5 and 6 and H3K27 monomethylation (a modification associated with chromatin condensation) (35,36) by H3.3 and H3K27me3 in *ddm1*, and the subsequent reincorporation of H3.1 in the absence of CLF. In this respect, the *ddm1 clf* mutant represents a valuable system to test the causal and functional relationship between DNA methylation and DNA compaction, which is elusive. As for the possibility of a direct antagonism between H3K27me3 and DNA methylation, we do not exclude it but the use of PcG mutants to analyze the impact of H3K27me3 loss at specific loci currently makes it difficult to tear apart cis-effects of this loss on a given TE from global and indirect effects. In the future, targeted removal of H3K27me3 by epigenome-editing needs to be implemented for this purpose.

Finally, our work in *ddm1 clf* mutant could have important implications for biological situations where TEs are naturally hypomethylated, DDM1 absent and/or chromatin decompacted. First, in the pollen vegetative nucleus, where hundreds of TEs are DNA-demethylated due to the activity of DEMETER glycosylases (37), where DDM1 is not detected (38). TE activation in this cell type is contributed by chromatin decondensation and DNA demethylation-dependent and independent mechanisms (39) and has been proposed to reinforce silencing in the gamete through production of small RNAs that could transit into the sperm cell (37,38,41,42). Whether PRC2 targets some of the demethylated TE-sequences and whether this could antagonize chromatin decondensation and modulate TE activation and small RNA production in this context is an open question. In this scenario, the mechanisms we have described in this work could play an important role in the reinforcement of silencing in the gametes and its modulation. Similarly, in the endosperm, the nutritive terminal tissue surrounding the seed and derived from the central cell, the chromatin is less condensed than in other types of nuclei (43). In addition, hundreds of TEs of the maternal genome are naturally hypomethylated forming primary imprints and many of these hypomethylated TEs are targeted by PRC2 forming secondary imprints (44,45). These secondary imprints cause, for instance, the silencing of the maternal *PHERES* locus while the cognate paternal allele is activated (45,46). The interplay between DNA methylation and PRC2 that we have evidenced could thus be particularly relevant in this cell type and modulate the imprinting of some genes as previously suggested (47). Finally, an exciting possibility is that PRC2 could target transposons after their mobilization and integration and this could slow down the establishment of their DNA methylation-based transgenerational and stable epigenetic silencing.

## Material and methods

### Plant material and growth condition

14 day-old seedlings grown on MS plates and with an 8h light/16h dark photoperiod were used throughout the study except for Southern Blot where analyses were performed on 5-week-old rosette leaves grown in the same conditions.

### Mutant lines

We used the *ddm1-2* allele (48) and the *clf-29* allele (49). Double *ddm1 clf* mutants were generated by crossing the above-mentioned mutants (19). Experiments were performed on F3 progenies using *ddm1*-2 2_nd_ generation plants as controls. We refer to the different progenies tested as different *ddm1-clf* lines or Biological Replicates (BR) thoughout the study.

### SDS-PAGE and Western blotting

Chromatin-enriched protein fraction was extracted from 10-days-old seedlings as described in (50), quantified by standard BCA assay and 20 µg were resolved on SDS/PAGE. After electroblotting the proteins on a Polyvinylidene difluoride (PVDF) membrane, H3K27me3 analysis was performed using an antibody against the H3K27me3 (Millipore, 07-449) at a 1:1000 dilution and a secondary antibody against rabbit coupled to HRP (Promega W4011) at a 1:20 000 dilution.

### BS-seq libraries

DNA was extracted using a standard CTAB protocol. Bisulfite conversion, BS-seq libraries and sequencing (paired-end 100 nt reads) were performed by BGI Tech Solutions (Hong Kong). Adapter and low-quality sequences were trimmed using Trimming Galore v0.3.3. Mapping was performed on TAIR10 genome annotation using Bismark v0.14.2 (51) and the parameters: -- bowtie2, -N 1, -p 3 (alignment); --ignore 5 --ignore_r2 5 --ignore_3prime_r2 1 (methylation extractor). Only uniquely mapping reads were retained. The methylKit package v0.9.4 (52) was used to calculate differential methylation in 100 bp or 20bp non-overlapping windows (DMRs). Significance of calculated differences was determined using Fisher’s exact test and Benjamin-Hochberg (BH) adjustment of p-values (FDR<0.05) and methylation difference cutoffs of 40% for CG, 20% for CHG and 20% for CHH. Differentially methylated windows within 100bp or 20bp of each other were merged to form larger DMRs. 100bp windows with at least six cytosines covered by a minimum of 6 (CG and CHG) and 10 (CHH) reads in all libraries were considered.

### RNA extraction and 3’quant-seq (RNA) analyses

Total RNA was extracted using Trizol followed by clean-up on RNeasy Plant Mini kit columns (Macherey-Nagel). 3’end librairies were prepared using the QuantSeq 3’ Fwd library prep kit (Lexogen). Libraries were sequenced to acquire 150 bp-reads on a NextSeq Mid output flow cell (Fasteris, Geneva). Expression level was calculated by mapping reads using STAR v2.5.3a (53) on the *A. thaliana* reference genome (TAIR10) with the following arguments -- outFilterMultimapNmax 50 --outFilterMatchNmin 30 --alignSJoverhangMin 3 -- alignIntronMax 10000. Counts were normalized and annotations (genes and TEs) were declared differentially expressed between samples (mutants vers wild-type) using DESeq2 (54).

### Chromatin immunoprecipitation and ChIP-seq analyses

ChIP experiments were conducted in two to four biological replicates of WT, *ddm1, clf* and *ddm1 clf* using an anti-H3K27me3 antibody (Millipore, 07-449). For each biological replicate, two IPs were carried out using 80 μg of Arabidopsis chromatin mixed with 5 μg of Drosophila chromatin, as quantified using BiCinchoninic Acid assay (Thermo Fisher Scientific). DNA eluted and purified from the two technical replicates was pooled before library preparation (Illumina TruSeq ChIP) and sequencing (Illumina sequencing single-reads, 1 × 50 bp) of the resulting input and IP samples performed by Fasteris (Geneva, Switzerland).

Reads were mapped using Bowtie2 v2.3.2 (55) onto TAIR10 *Arabidopsis thaliana* and Drosophila melanogaster (dm6) genomes. Normalization factor (Rx) for each sample using spiked-in *D. melanogaster* chromatin was calculated (56) using the following formula Rx = r/Nd_IP, where Nd_IP corresponds to the number of reads mapped on *D. melanogaster* genome in the IP and r corresponds to the percentage of *Drosophila-*derived reads in the input. Genomic regions significantly marked by H3K27me3 were identify using MACS2 (57) and genes or TEs overlapping these regions were obtained using bedtools (58). The number of reads over marked genes or TEs were normalized by applying the normalization factor and differentially marked genes between samples were calculated using DESeq2 (54).

### TE-sequence capture

TE sequence capture was performed on around 100 seedlings in all cases except *ddm1-clf*#23 NOchl were only 12 seedlings were recovered. Seedlings were grown in plates under control (long-day) conditions and genomic DNA was extracted using the CTAB method (59). Libraries were prepared as previously described (27) using 1μg of DNA and TruSeq paired-end kit (Illumina) following manufacturer instructions. Libraries were then amplified through 7 cycles of ligation-mediated PCR using the KAPA HiFi Hot Start Ready Mix and primers AATGATACGGCGACCACCGAGA and CAAGCAGAAGACGGCATACGAG at a final concentration of 2µM. 1μg of multiplexed libraries were then subjected to TE-sequence capture exactly as previously reported (29). Pair-end sequencing was performed using one lane of Illumina NextSeq500 and 75bp reads. About 42 million pairs were sequenced per library and mapped to the TAIR10 reference genome using Bowtie2 v2.3.2 (55) with the arguments --mp 13 --rdg 8,5 --rfg 8,5 --very-sensitive. An improved version of SPLITREADER (available at https://github.com/LeanQ/SPLITREADER) was used to detect new TE insertions. Briefly, split-reads as well as discordant reads mapping partially on reference and consensus TE sequences (obtained from RepBase update) were identified, soft clipped and remapped to the TAIR10 reference genome using Bowtie2 (55). Putative insertions supported by at least one split- and/or discordant-reads at each side of the insertion sites were retained. Insertions spanning centromeric repeats or coordinates spanning the corresponding donor TE sequence were excluded. In addition, putative TE insertions detected in more than one library were excluded to retain only sample-specific TE insertions.

### Southern blot

DNA from 5 week-old rosette leaves was extracted using a standard CTAB protocol. 1,5 μg of genomic DNA was digested overnight with SSpI restriction enzyme. The digestion was run on a 1% agarose gel, transferred to Hybond N+ membranes, blocked and washed according to manufacturer instructions (GE Healthcare). Membranes were probed with a PCR product (corresponding to a fragment of *EVD* GAG sequence,) radiolabeled with alpha 32P-dCTP using the Megaprime DNA Labeling System. EVD PCR product was generated with the same primers as in (60).

### Pyrosequencing

*ATCOPIA93* DNA from gDNA (same extraction as used in Southern Blot) was analyzed as in (19).

### Cytology

DAPI staining on fixed nuclei was performed as described in Bourbousse et al., 2015 using 10-days-old chopped cotyledons to avoid developmental defects due to accumulation of transposition. For immuno-detection, a primary antibody against H3K9me2 (Abcam 1220) at a 1:200dilution was used, followed by a secondary antibody (Alexa Fluor 488 mouse) at a 1:400 dilution and DAPI staining. Images were acquired with a confocal laser scanning microscope and processed using ImageJ (https://imagej.nih.gov/ij/).

### Data availability

Sequencing data has been deposited in the European Nucleotide Archive (ENA) under project PRJEB34363.

## Acknowledgments

We thank the Genomics and Informatics facilities at IBENS as well as the Barneche lab (C. Bourbousse and G. Teano) for their help with Drosophila spike-in ChIP experiments and discussions. This work was funded by a Human Frontier Scientific Program Career Development Award (HFSP-CDA-00018/2014) granted to A.D and the Centre National de la Recherche Scientifique (CNRS) Momentum program granted to L.Q. Additional support was from the CNRS.

## Conflict of interest

The authors declare that they have no conflict of interest.

---

**Sup Fig.S1A.**
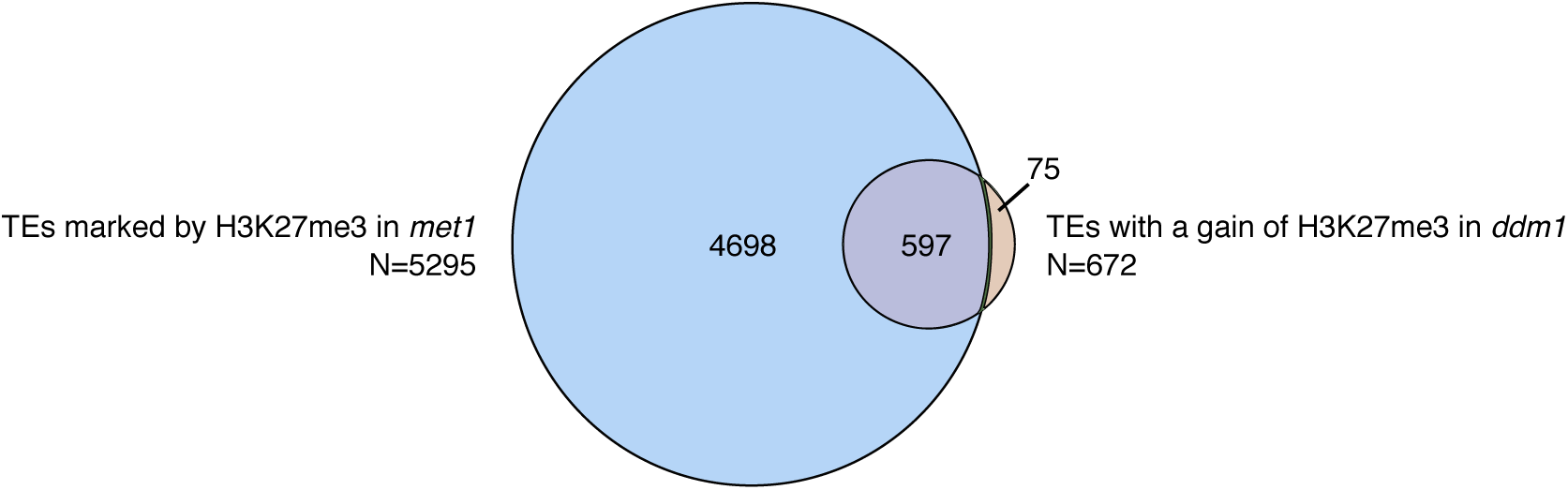
Overlap between TEs marked by H3K27me3 in *met1* (Deleris et al., 2012) and TEs with a gain of H3K27me3 in *ddm1*.

**Sup Fig.S1B.**
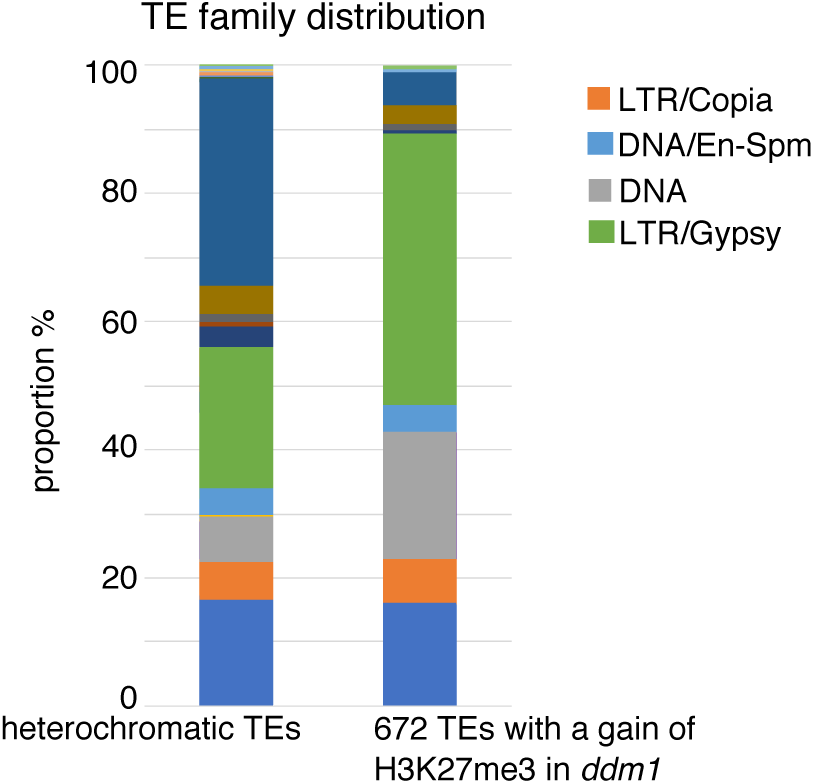
Distribution of the different TE families among the TEs that significantly gain H3K27me3 in *ddm1* compared to the family distribution of heterochromatic, pericentromeric TEs (targets of DDM1).

**Sup Fig.S1C.**
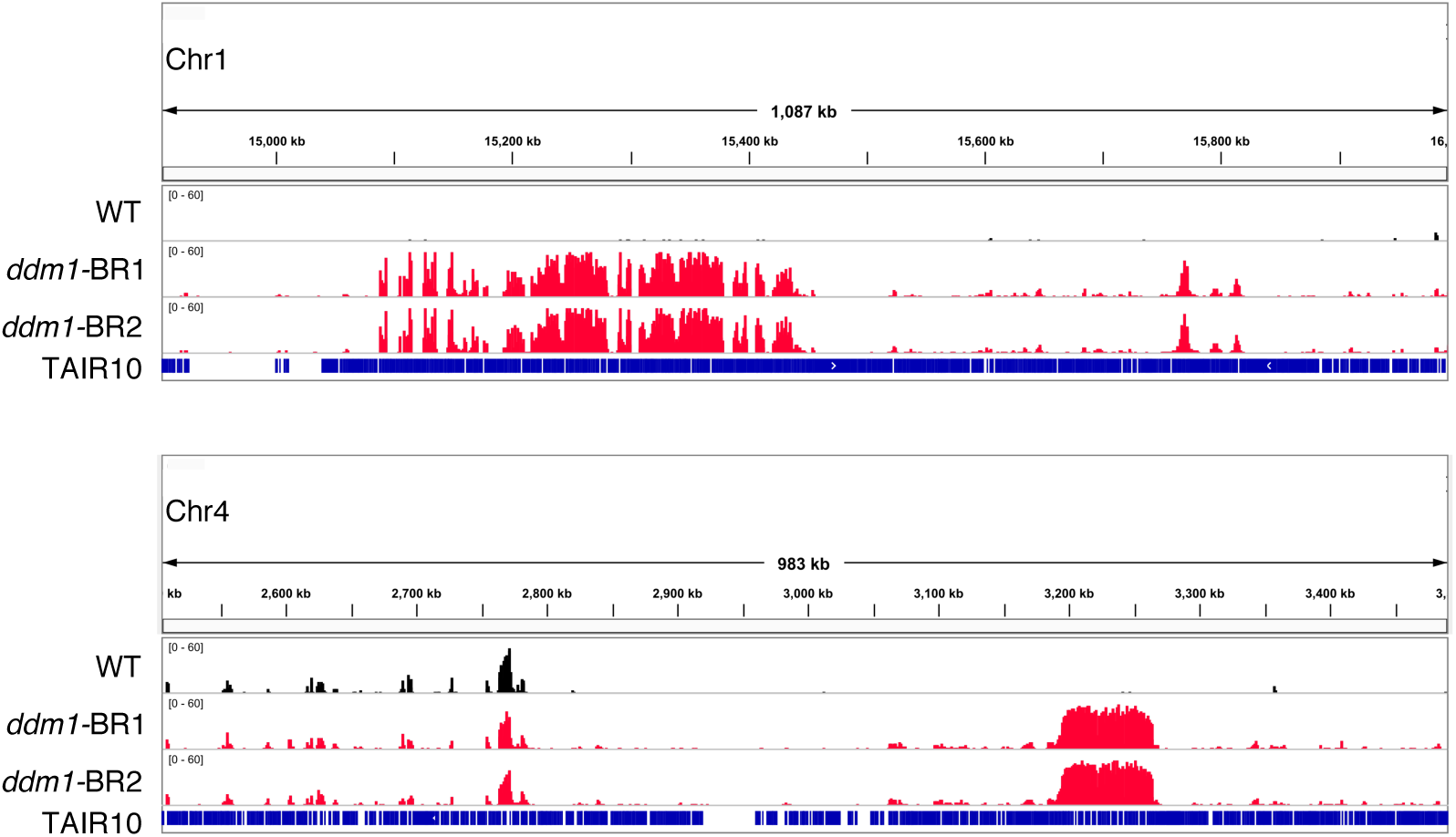
Genome browser view of chromosomes 1 and 4 showing the two largest H3K27me3 domains in *ddm1* (BR=biological replicate).

**Sup Fig.S2A.**
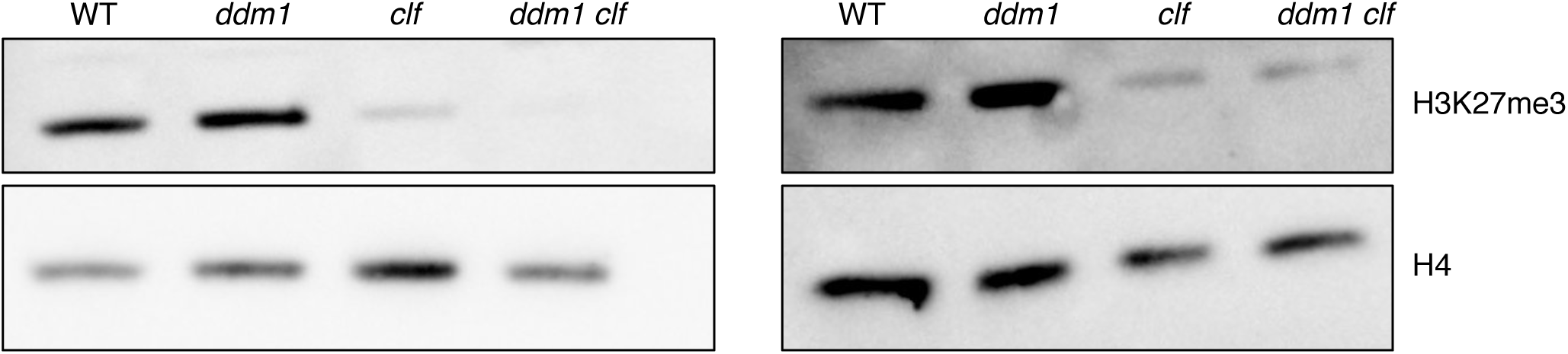
Immuno detection of H3K27me3 in chromatin extracts (H4 is used as a loading control), two biological replicates are shown.

**Sup Fig.S2B.**
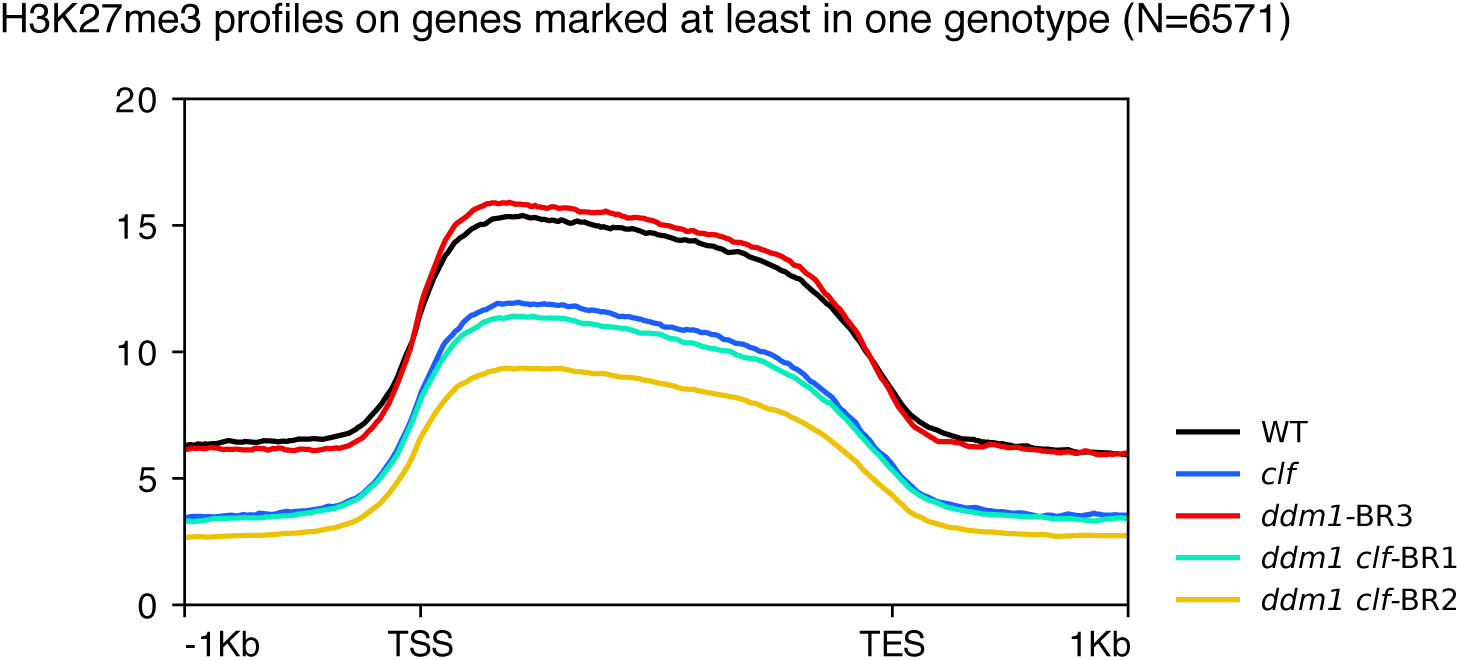
Metagene showing H3K27me3 levels on genes marked in at least one genotype

**Sup Fig.S2C.**
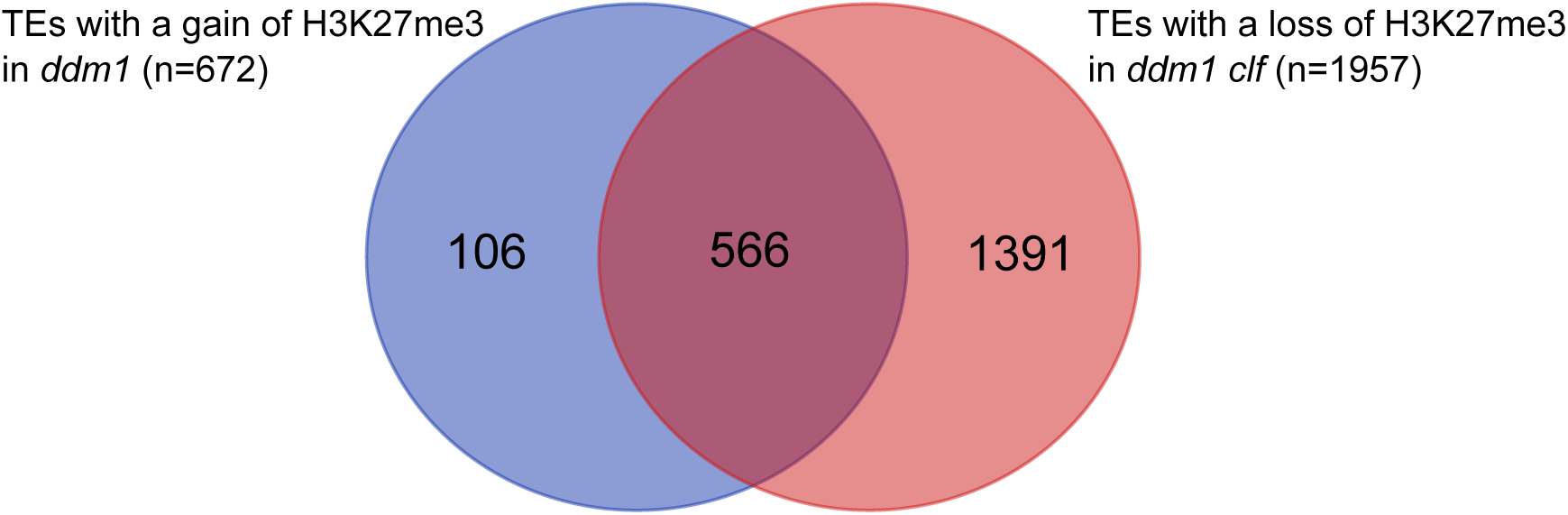
Venn diagram showing the overlap between the TEs that gain H3K27me3 in *ddm1* and the TEs that lose H3K27me3 in *ddm1 clf*.

**Sup Fig.S3A.**
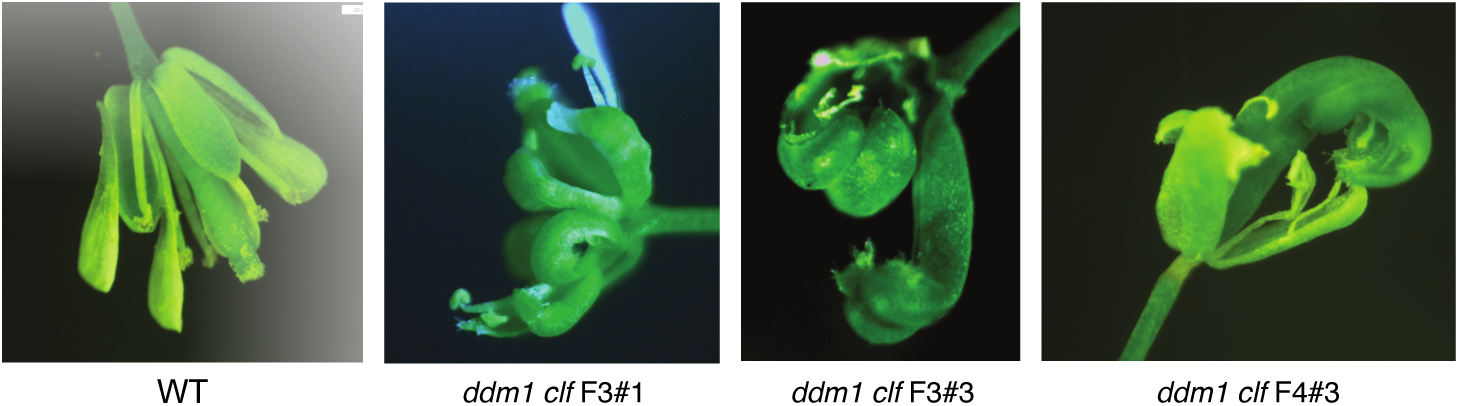
Representative pictures of *ddm1 clf* mutant flowers coming from different F3 (used for the southern blot) or F4 progenies alongside WT flowers.

**Sup Fig.S3B.**
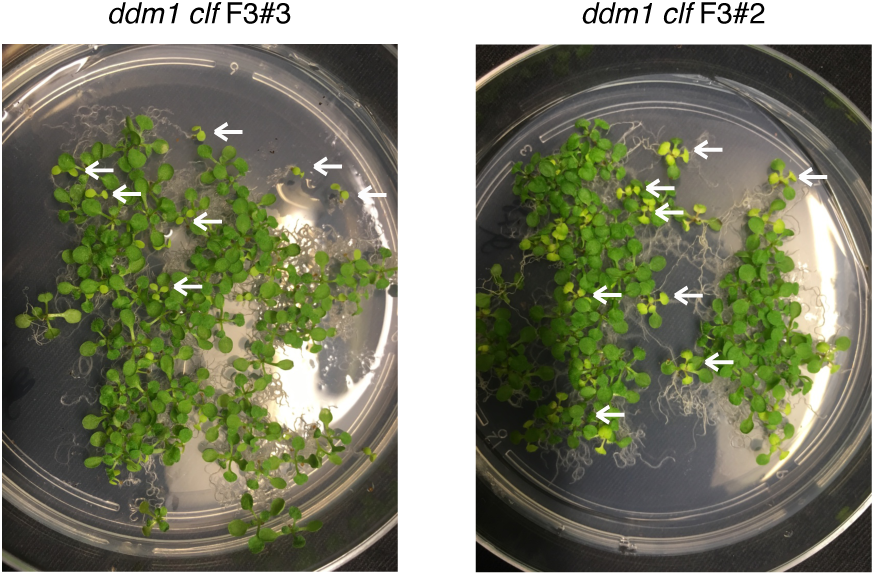
Representative picture of 15 days-old *ddm1 clf* F3 seedlings coming from two independent F2 lines; the white arrows show two different phenotypes segregating 1:3, growth arrest (left) and chlorosis (right) respectively.

**Sup Fig.S3C.**
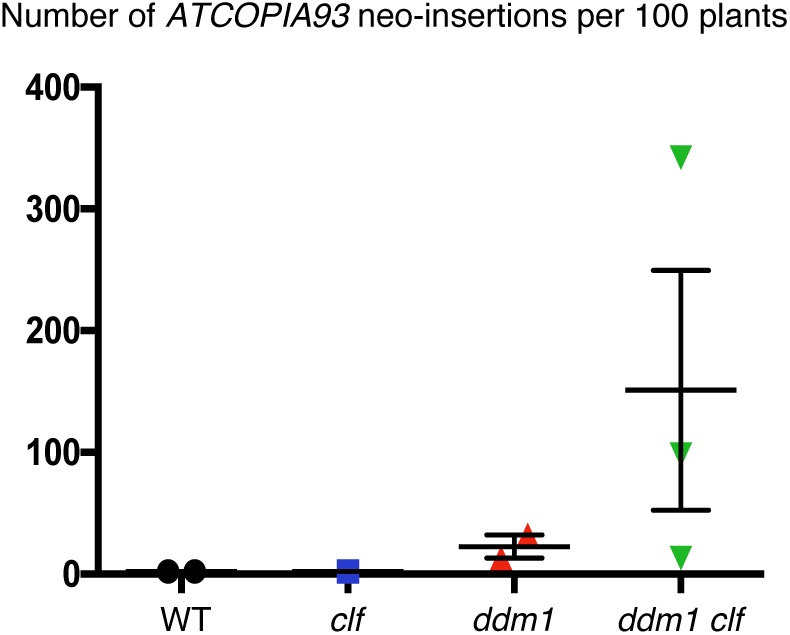
Graph showing the number of *EVD* neo-insertions revealed by TE-capture in lines derived from a cross between *ddm1* and *clf* mutants.

**Sup Fig.S3D.**
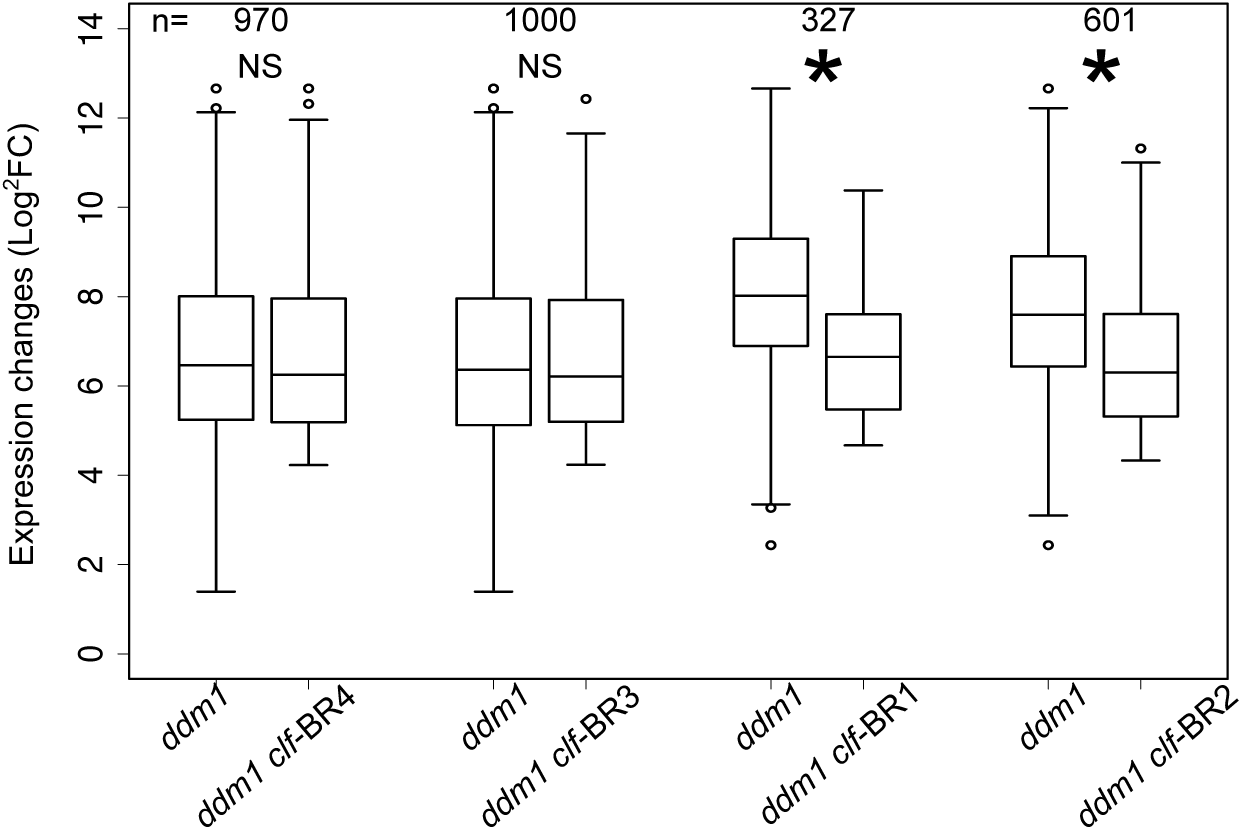
Box plot of log2 RNA fold changes (mutant/WT) over the TEs that are transcriptionally active in *ddm1 clf*. * indicates a significant decrease in the fold change for a given *ddm1 clf* line relative to *ddm1* (Pval<0,05 t-test)

**Sup Fig.S5A.**
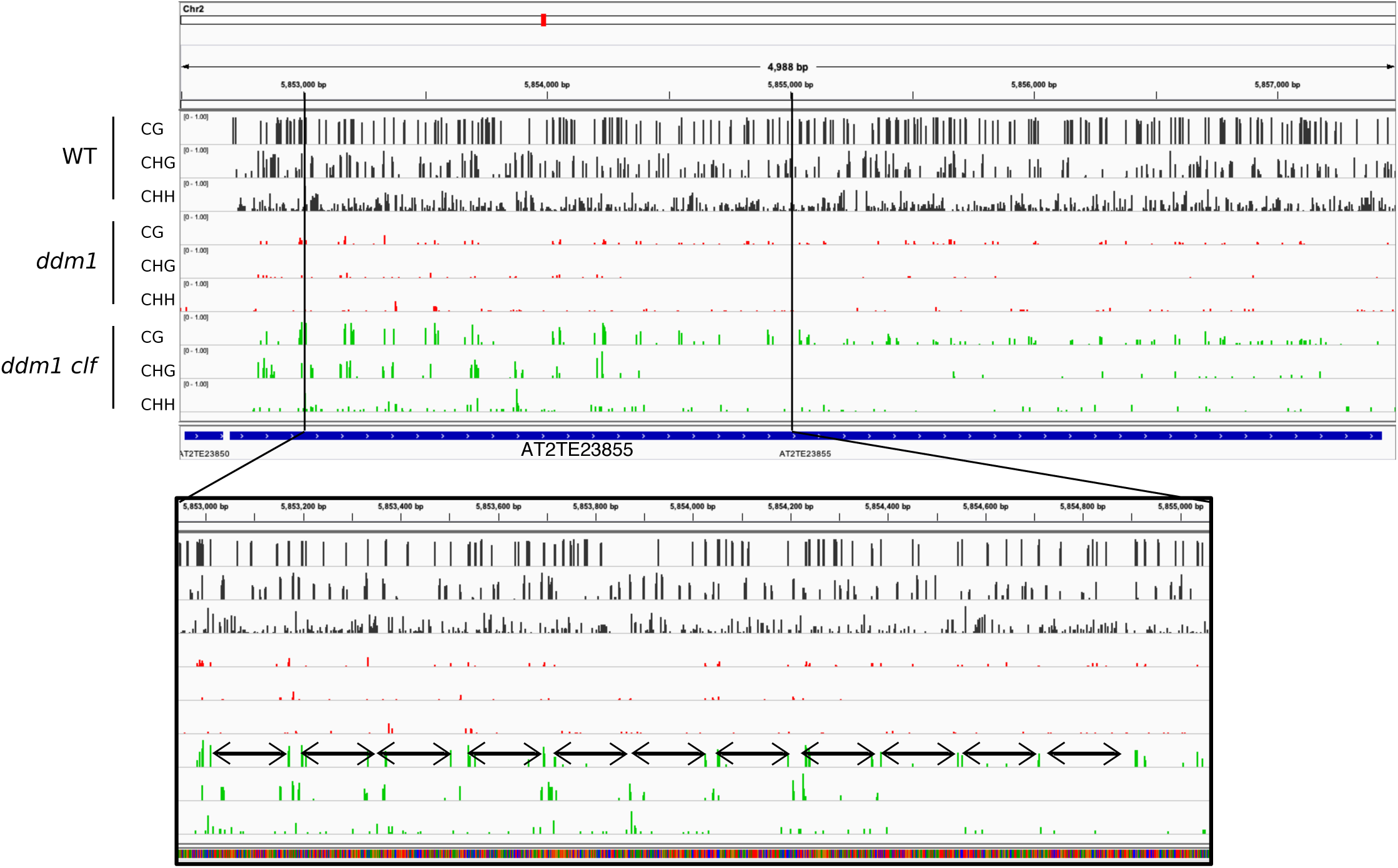
Representative view of a TE with a periodic DNA hypermethylation every 140bp; the bottom panel is a zoom of the top panel and the double arrow represents a distance of 140bp.

**Sup Fig.S5B.**
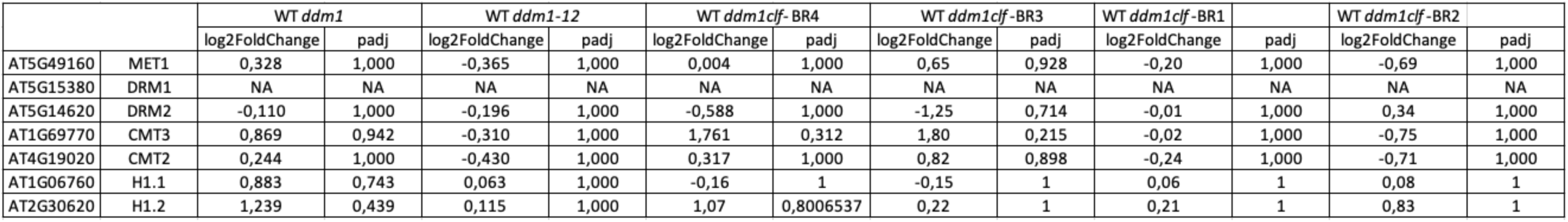
Table showing expression changes of genes involved in DNA methylation in different *ddm1* and *ddm1 clf* populations analysed in this study.

## Notes

### Competing Interest Statement

The authors have declared no competing interest.

